# *In silico* modeling and interactive profiling of BPH resistant R genes with elicitor molecules of rice planthoppers

**DOI:** 10.1101/2021.10.25.465708

**Authors:** Krishnamanikumar Premachandran, Thanga Suja Srinivasan

**Affiliations:** Centre for Climate Change Studies, International Research Centre, Sathyabama Institute of Science and Technology, Chennai 600119, Tamil Nadu, India

**Keywords:** Brown planthopper, White backed planthopper, Small brown planthopper, BPH resistant genes, Insect salivary secretome, NBS-LRR region, Protein-protein interaction

## Abstract

Brown planthopper resistant NBS-LRR specific R genes (*Bph9, Bph14, Bph18, Bph26*) have been reported in rice. BPH specific R genes were clustered with other R genes of rice on chromosome 12 (*Bph9, Bph18, Bph26)* and 3 (*Bph14*). Motif analysis of BPH specific R genes showed the predominant motifs as CC, NBS and LRR regions. *Bph9, Bph18* and *Bph26* R genes exhibited high degree of sequence similarity in their CC and NBS region and are considered as functional alleles of BPH resistance at chromosome 12. LRR region of BPH genes were interacting with the elicitor molecules of planthoppers and are the potential lignad binding site. *Bph14* exhibited more number of LRR repeats and were interacting efficiently with all the tested salivary elictor molecules of planthoppers. *Bph18* with no LRR region exhibited reduced interaction efficiency with the tested elicitor molecules of planthoppers. Our *in silico* studies confirms that *Bph14 R* gene resistance protein to be a promising candidate for providing broad spectrum resistance against planthoppers of rice. The study further provides new avenues to investigate the mechanism of receptor-ligand recognition and signaling mechanism against rice planthoppers.

## Introduction

Rice planthoppers (brown planthopper (BPH), white backed planthopper (WBPH) and small brown planthopper (SBPH)) are major phloem feeding insects causing severe yield loss every year in major rice growing regions of the world. Recent outbreaks of rice planthoppers (BPH, WBPH and SBPH) have been reported in China (2005-2006) [1]; Indonesia (2009), Vietnam (2010) [2] and Northern India (2013-2015) [3]. Since the early 1970s, rice varieties have been bred for resistance to plant- and leafhoppers especially *Nilaparvata lugens* (brown planthopper). Breeding for planthopper resistance has resulted in the identification of nearly 39 resistant gene loci/QTLs against BPH, 8 againt WBPH and 3 againt SBPH from wild and cultivated rice varieties and landraces. A few of these resistant gene loci (i.e. *Bph1, bph2, Bph3, bph8* and *Bph9*) have been successfully introgressed into modern rice varieties, however insect biotypes/populations emerge that overcome these plant traits within a few years of varietal deployment and the mechanism of planthopper adaption to resistant rice varieties is still unclear.

During the course of evolution, plants have developed an innate immune system to respond against the attackers. R genes are resistant genes present in the plant genome responsible for plant disease resistance. They are members of plant immune receptors encoding proteins that specifically recognize pathogen associated virulence factors and provoke plant immunity. R genes mostly encode NBS-LRR protein with a common modular structure comprising of amino terminal nucleotide binding site (NBS) and C-terminal leucine rich repeats (LRR) region [4]. The resistant (R) genes of plants directly or indirectly interacts with effector protein therefore sensing pathogen attack and inducing disease resistance [5]. More than 400 R genes have been reported so far in rice plants [6] and 6 R genes have been reported against BPH insect [7]. Nucleotide Binding Site-Leucine Rich Repeat (NBS–LRR) proteins have been identified in BPH resistance loci (*Bph14, Bph18, Bph9, Bph26*) where they function similar to receptors for pathogen detection, linking plant innate immunity to planthopper resistance [8]. The central nucleotide-binding domain of NBS-LRR controls the ATP/ADP-bound state mediating downstream signaling and the N-terminal coiled-coil (CC) or Toll/Interleukin-1 receptor (TIR) domains are used as signaling either cellular targets of effector action or with downstream signaling components [9]. The C-terminal LRR domains of NBS-LRR proteins are variable in length and forms series of β-sheets that interact with the effector molecules [10].

Recent studies on the salivary components of rice planthoppers provide evidence for their role in successful rapid adaptation to host immunity genes [11]. Herbivore salivary components can modify the innate defense mechanism of host plants aiding in better protection, nutrient access and adaptation. Planthoppers secreates gelly and watery saliva from the salivary glands which aid in insect feeding [12]. The watery saliva contains hydrolyzing, digestive and cell wall degrading enzymes. Gelly saliva used to form a continuous salivary sheath providing mechanical stability,lubrication and protection against plant defense chemicals for the insect [12]. Salivary proteins were reported to act as a effectors/elicitors interacting with host resistant genes and activating complex defence responses like mitogen-activated protein kinase (MAPK) cascades, reactive oxygen species (ROS), jasmonic acid (JA), salicylic acid (SA), and ethylene(ET) signaling pathways [13].

In rice planthoppers, no prior studies have been conducted to investigate the interaction of BPH resistant R genes and salivary elicitors of rice planthoppers. The proposed study provides a framework about the structure and interaction of different characterized BPH resistant R genes with salivary elicitors of rice planthoppers. The sequential and structural similarity among characterized BPH resistant genes of rice were carried out and their domains and binding sites were identified. The *in silico* study provide new insights into the structural interaction of BPH specific NBS LRR genes with insect specific elicitor molecules. We used few other R genes of rice like blight and blast associated genes of rice to validate the structural similarlity, and functional mimicry among R genes of rice. Our results can assist in further understanding the interaction between R protein-elicitor perception and plant defense signaling against planthoppers of rice. The study can form a baseline for further exploring salivary elicitor mediated adaptation and evolution biology against plant resistant genes

## Materials and methods

Thirty nine major BPH resistant genes have been identified so far (Zhang *et al* 2020). Of which four well characterized BPH resistant genes (*Bph9, Bph14, Bph18* and *Bph26*) possessing NBS-LRR region and ten different salivary elicitors (carboxylesterase, cathepsin L, chitinase like protein, serine protease, aminopeptidase N, cathepsin B like protease, dipeptidyl peptidase, phosphoglycerate kinase 1, carboxylesterase and serine protease) of rice planthoppers (BPH, WBPH and SBPH) have been taken for the study. Other R genes of rice *Xa1, Xa14* (bacterial blight (*Xanthomonas oryzae pv. oryzae*)) *Pib* and *Pid3 (*blast (*Magnaporthe oryzae*)) were also taken for the study. Protein sequences were obtained from the UniProt protein database (https://www.uniprot.org/) [15], a freely available open access protein databank and MEROPS the peptidase database (https://www.ebi.ac.uk/merops/) [16].

### Multiple sequence alignment

Multiple sequence alignment (MSA) among four different BPH resistant genes and other R genes of rice (*Xa1, Xa14, Pib* and *Pid3*) were carried out to identify the sequence similarities, conserved sites and evolution among BPH genes and other R genes of rice. MSA were carried out using MEGA (Molecular Evolutionary Genetics Analysis) software, version 10 [17]. BPH and other resistant R gene (*Xa1, Xa14, Pib* and *Pid3*) sequences were aligned by ClustalW multiple sequence alignment program with default parameter settings using MEGA software.

### Phylogenetic analysis

Unrooted phylogenetic tree was constructed using maximum likelihood method to analyse the evolutionary relationship among four BPH genes and other R genes of rice. Phylogenetic tree was constructed by MEGA (Molecular Evolutionary Genetics Analysis) software, version 10 [17] and the reliability was checked by setting up the bootstrap replication value as 1000. Other default parameters remain same.

### Physio chemical analysis of BPH resistant genes

Physical and chemical properties of BPH resistant proteins were analyzed computationally with ProtParam tool (https://web.expasy.org/protparam/) [18]. Various physical and chemical properties such as molecular weight, theoretical pI, amino acid composition, extinction coefficient, instability index, aliphatic index and grand average of hydropathicity (GRAVY) were calculated.

### Motif identification

Conserved motifs were identified by MEME (Multiple Em for Motif Elicitation) software, version 5.1.1) [19] among BPH and other R genes of rice and minimum width of the motif was set as 6 amino acids and the maximum width was set as 50 amino acids and other parameters remained same as default.

### NBS-LRR region prediction

Coiled coil (CC) region of all BPH resistant genes were predicted with COILS server (https://embnet.vital-it.ch/software/COILS_form.html) (Lupas et al., 1991). Nucleotide Binding Site (NBS) region and Leucine Rich Repeats (LRR) region of *Bph9, Bph14, Bph18* and *Bph26* proteins were predicted by using Pfam (http://pfam.xfam.org/) [21] and InterPro (https://www.ebi.ac.uk/interpro/), an online webserver for functional analysis of protein and identification of important domain and sites [22].

### Preliminary analysis of protein-protein interaction sites by iLoops

iLoops tool was used to predict the interaction between BPH resistant proteins and insect salivary proteins (hereafter called as elicitor molecule) Galaxy InteractoMIX iLoops (http://galaxy.interactomix.com) [23] version 0.1, an integrated computational platform was used to predict the BPH resistant protein (*Bph9, Bph14, Bph18* and *Bph26*) and insect salivary elicitors (carboxylesterase, cathepsin L, chitinase like protein, serine protease, aminopeptidase N, cathepsin B like protease, dipeptidyl peptidase, phosphoglycerate kinase 1, carboxylesterase and serine protease) interaction sites by identification of similar structural loops, conserved motifs and interactive domains based on ArchDB (structural classification of loops in proteins) and SCOP (structural loop database).

### Homology modelling

BPH resistant protein and insect salivary elicitor molecule 3D structures were developed by Swiss model web server (https://swissmodel.expasy.org/), a fully automated homology-based protein structure modelling server [24]. Templates were selected for modelling the BPH and other insect salivary elicitors by BLAST program on Protein Data Bank and selected based on the maximum sequence identity. Selected templates were used to model the structures and further minimization of the predicted structures were carried out by using SWISS-PDB Viewer (SPDBV) software, version 4.10 [25]. Ramachandran plot was plotted with PROCHECK webserver (https://www.ebi.ac.uk/thornton-srv/software/PROCHECK/) for structural conformation, stereochemical quality validation of the predicted 3D protein models of BPH resistant proteins [26].

### Protein-protein docking and analysis

All of the four BPH resistant proteins were docked with each ten insect salivary elicitors individually in ClusPro Protein-Protein docking web server (https://cluspro.bu.edu/) [27] version 2.0 and the docked models were selected based on the appropriate interactive site and docking score. PDBsum web server (https://www.ebi.ac.uk/thornton-srv/databases/pdbsum/Generate.html) [28] was used to analyse the interaction (number of hydrogen bonds, non-contacted interaction and salt bridges etc.,) between BPH resistant proteins and insect salivary elicitors. The residue interactions across the interface and the distance between the two interacting amino acid residues were analysed. All the analysis of the structured models was carried out by using PyMOL molecular visualization software, version 2.4.0 and BIOVIA Discovery studio visualizer software, version 20.1.0 [29], [30].

## Results

### Sequence analysis

Protein sequences of BPH resistant genes and insect salivary elicitor molecules of BPH, WBPH and SBPH were obtained from databases and subjected for sequence analysis. Multiple sequence alignment shows that *Bph9, Bph18* and *Bph26* gene have significant similarity whereas *Bph14* shows less similarities with other BPH resistant proteins and higher similarity with *Xa1* and *Xa14* gene. *Bph14* exhibits higher similarity with *Xa1* and *Xa14* in the NBS region and 6 conserved LRR repeats observed between *Xa1, Xa14* and *Bph14* (Data not shown). Further, *Bph14* has a smaller LRR region compared to *Xa1* and *Xa14* genes (Data not shown). Rice blast resistant genes (*Pib* and *Pid3*) shows higher similarity with *Bph9* gene than other genes. Amino acid length of *Bph18* protein was comparatively less than other BPH resistant proteins but showed 94%, 87.5% and 11.5% similarity with *Bph26, Bph9* and *Bph14* respectively. *Bph9* protein shows 87.5%, 84% and 12% sequence similarity with *Bph18, Bph26* and *Bph14* respectively. *Bph14* shows 11% and 12% aminoacid sequence similarity with *Bph18* and *Bph26* accordingly and 15% similarity with both *Xa1* and *Xa14* gene. *Xa1* and *Xa14* gene shows 84% sequence similarity within them. *Pib* gene shows about 40% similarity with all BPH resistant genes except *Bph14* and *Pid3* shows very minimal sequence similarity with BPH resistant genes and other blight resistant genes.

Unrooted phylogenetic analysis of BPH resistant genes with other R genes of rice revealed the evolutionary relationship within BPH resistant R genes and other R genes of rice.We observed two different clades; clade I includes *Bph9, Bph18, Bph26, Pib* and *Pid3* genes and clade II includes *Bph14, Xa1* and *Xa14* (Fig 2). Interestingly *Bph14* gene comes under clade II with the resistant R genes of rice (*Xa1* and *Xa14*) with a bootstrap value of 100. Bootstrap values were 100% for all except *Bph18*/*Bph26* (95%) in clade I (Fig 2). MSME software has been used to identify the significant motifs and motif arrangements of BPH resistant proteins and other R genes of rice. Twenty different motifs were obtained. *Bph9* and *Bph26* shares exactly same number of motifs and motif arrangements along with *Pib* (Fig 3). *Bph18* shows much similar with *Bph9/Bph26* motif arrangements nevertheless number of motifs were less. The number and arrangement of motifs identified in *Bph14* varied when compared with other BPH R genes. *Xa1* and *Xa14* shows identical number of motifs and arrangements (Fig 3).

### Physio chemical analysis of BPH resistant genes

Physiochemical properties of BPH resistant proteins were predicted computationally and given in Table 2. Molecular weight of *Bph9, Bph14* and *Bph26* predicted to be 136.7, 149.1 and 138.4 kDa in size respectively and *Bph18* 77.2 kDa in size. Isoelectric point of *Bph9, Bph14, Bph18* and *Bph26* predicted to be 8.3, 6.1, 7.5 and 8.7 accordingly (Table 2). All the proteins consist equal amount of positively charged and negatively charged amino acid residues. Instability values predicted to be above 40 for all the proteins with above 90 aliphatic index and GRAVY values were -0.282, -0.263, -0.327 and -0.278 for *Bph9, Bph14 Bph18* and *Bph26* respectively (Table 2).

### NBS-LRR region prediction

Coiled coil region of *Bph9, Bph14, Bph18* and *Bph26* proteins were predicted to be present in 107-127, 32-66, 107-127 and 107-127 amino acid position of N terminal region of the proteins respectively (Fig 1A, Table 1). Pfam database and InterPro protein families and domains database was used to identify the nucleotide binding site and leucine rich repeats of the BPH proteins. *Bph9, Bph18* and *Bph26* proteins have duplicated NBS region while other BPH resistant protein have a single NBS region within the sequence (Fig 1C, Table 1). *Bph9* protein have NBS region at 159-352 amino acid followed by 405-728 and *Bph18* have NBS region at 158-351 and 404-680^th^ amino acid. *Bph26* predicted to have duplicated NBS region at the position of 158-366 and 419-742. *Bph14* have a single NBS region at 191-518^th^ amino acid position as well (Fig 1A, Table 1). On the other hand, *Bph9* protein have LRR region from 742-1160^th^ amino acid position where *Bph14* and *Bph26* proteins have LRR region at 537-1299 and 788-1209^th^ amino acid position respectively (Table 1). *Bph18* has not shown any LRR region in InterPro analysis. *Bph9* and *Bph26* protein has 10 LRR repeats whereas *Bph14* consists of 16 LRR repeats (Fig 1B, Table 1).

**Table 1.** Details of BPH resistant R genes *(Bph9, Bph14, Bph18* and *Bph26)* and their chromosomal location along with CC-NBS-LRR region details were given below. *Bph9, Bph18* and *Bph26* exhibited duplicated NBS region. *Bph18* doesn’t have LRR region.

| Gene | Chromosome location | NBS protein | CC region | NBS region | LRR region | LRR repeats |
| --- | --- | --- | --- | --- | --- | --- |
| *Bph9* | 12L | CC-NB-NB-LRR | 107-127 | 159-352, 405-728 | 742-1160 | 10 |
| *Bph14* | 3L | CC-NB-LRR | 32-66 | 191-518, | 537-1299 | 16 |
| *Bph18* | 12L | CC-NB-NB-LRR | 107-127 | 158-351, 404-680 | - | - |
| *Bph26* | 12L | CC-NB-LRR | 107-127 | 158-366, 419-742 | 788-1209 | 10 |

**Table 2.** Physio chemical properties of BPH resistant R genes *(Bph9, Bph14, Bph18* and *Bph26)* given below. Molecular weight, theoretical pI, amino acid composition, extinction coefficient (Ec^a^), instability index (II), aliphatic index (AI) and grand average of hydropathicity (GRAVY)

| S. No | Protein | Length | Molecular weight (kDa) | pI | (-) R | (+) R | Ec<sup>a</sup> | II | AI | GRAVY |
| --- | --- | --- | --- | --- | --- | --- | --- | --- | --- | --- |
| 1 | *Bph9* | 1206 | 136.7 | 8.34 | 159 | 167 | 142220 | 42.38 | 98.13 | -0.282 |
| 2 | *Bph14* | 1323 | 149.1 | 6.12 | 176 | 160 | 148720 | 48.99 | 93.70 | -0.263 |
| 3 | *Bph18* | 680 | 77.2 | 7.52 | 95 | 96 | 92080 | 43.97 | 91.90 | -0.327 |
| 4 | *Bph26* | 1218 | 138.4 | 8.74 | 156 | 172 | 144740 | 42.14 | 96.76 | -0.278 |

### Preliminary analysis of protein-protein interaction by iLoops

iLoops by galaxy InteractoMIX was used to scrutinize the list of salivary proteins (from BPH, WBPH and SBPH) that can interact with BPH resistant genes based on similar structural loops, conserved motifs and interactive domains. 46 salivary proteins were selected from BPH, WBPH and SBPH insect saliva and subjected for InteractoMIX. iLoops analysis showed 10 salivary proteins that have the interactive possibility with BPH resistant genes (Data not shown). Based on the preliminary interactive study using iLoops, 10 salivary elicitors were taken for further modelling and protein-protein docking interaction studies.

### 3D modelling

Amino acid sequence of BPH gene and insect salivary elicitors were retrieved from UniProt database and 3D structure were modelled by using comparative modelling method. Swiss-model was used to model the structure of BPH proteins and salivary elicitors of planthoppers. Based on the query coverage template was chosen and 3D model was executed. Disease resistance RPP13-like protein 4 from *Arabidopsis thaliana* was taken as template for model of all the BPH resistant proteins. BPH resistant proteins showed around 30% sequence similarity with RPP13-like protein 4 (Fig 4A, Table 3A). Ramachandran plot was used to validate the structural conformation of the predicted 3D structure of the proteins. More than 97% of the amino acid residues fall under most favorable region and additionally allowed region. Residues in disallowed regions were less than 1% (Fig 5). Carboxylic ester hydrolase (5ikx.1.A) and Cathepsin L1 (3hwn.1.A) was chosen as template for BPH salivary elicitors carboxylesterase and cathepsin L protease enzyme with 36.61% and 59.84% sequence similarity respectively (Fig 4B, Table 3B). Likewise, serine protease hepsin (5ce1.1.A) and insect group II chitinase (5y2b.1.A) was chosen as template for serine protease and chitinase like protein with sequence similarity of 36.05% and 30.98% accordingly. Aminopeptidase N, procathepsin B and phosphoglycerate kinase 1 from human were chosen as template model for aminopeptidase N, cathepsin B like protease and phosphoglycerate kinase 1 salivary protein of SBPH with sequence similarity of 26.84%, 43.55% and 63.35% respectively. Dipeptidyl phosphate 8 chosen as template model for dipeptidyl peptidase 4 enzyme from SBPH with 38.94% sequence similarity (Fig 4B, Table 3B). To model carboxylesterase and serine protease 6 enzymes from salivary protein of WBPH, carboxylic ester hydrolase and prostasin were selected as template model with 33% and 22.75% sequence similarity.

**Table 3A.** Comparative modelling 3D structure prediction of BPH resistant R proteins. Details of templates chosen for the structure prediction along with Q mean value, GMQE and query coverage were given.

| S. No | Protein name<br>(UniProt accession no) | Template<br>(PDB id) | Q mean | GMQE | Query<br>coverage |
| --- | --- | --- | --- | --- | --- |
| 1 | *Bph9* | Disease resistance<br>RPP13-like protein 4<br>(6j5t.1.C) | -6.17 | 0.14 | 32.34 % |
| 2 | *Bph14* | Disease resistance<br>RPP13-like protein 4<br>(6j5t.1.C) | -4.60 | 0.27 | 29.16 % |
| 3 | *Bph18* | Disease resistance<br>RPP13-like protein 4<br>(6j5t.1.C) | -2.72 | 0.18 | 32.51 % |
| 4 | *Bph26* | Disease resistance<br>RPP13-like protein 4<br>(6j5t.1.C) | -5.15 | 0.13 | 31.64 % |

**Table 3B.**
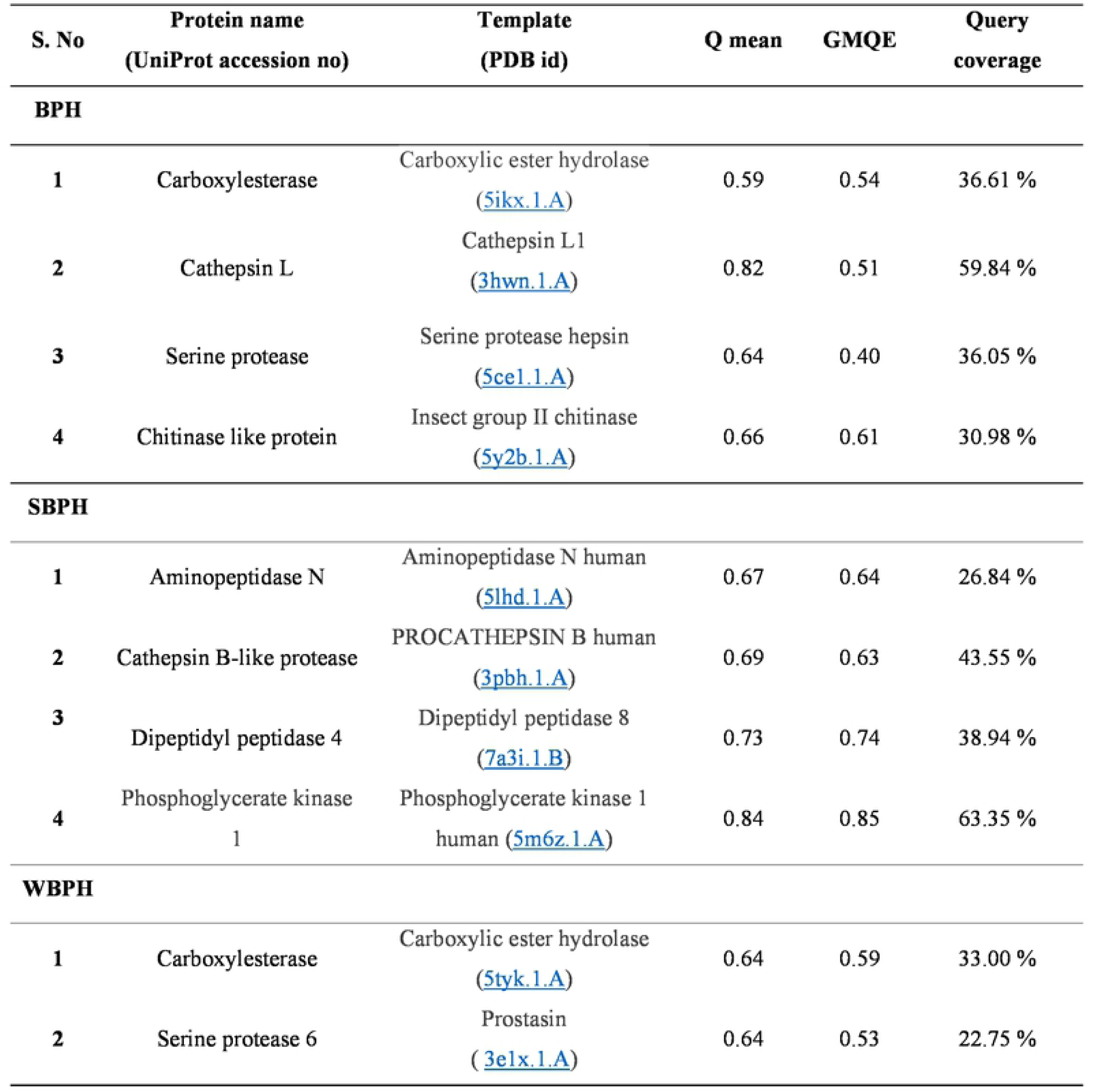
Comparative modelling 3D structure prediction of BPH insect salivary elicitors. Details of templates chosen for the structure prediction along with Q mean value, GMQE and query coverage were given.

### protein-protein interaction and docking

ClusPro protein-protein docking webserver used to dock BPH resistant genes with all ten salivary elicitors. *Bph14* protein showed higher interaction among all insect salivary elicitors followed by *Bph26, Bph9* and *Bph18* (Fig 6). *Bph26* protein shows upmost interaction with dipeptidyl peptidase 4 salivary protein of BPH with docking score of -1331.8 and 23 hydrogen bonds (Fig 6, Table 4). *Bph14*-dipeptidyl peptidase IV, *Bph14*-aminopeptidase N, *Bph26*-dipeptidyl peptidase IV, *Bph14*-serine protease, *Bph9*-dipeptidyl peptidase IV, *Bph14*-carboxylesterase, *Bph9*-serine protease, *Bph9*-aminopeptidase N, *Bph14*-cathepsin B and*Bph26*-aminopeptidase N models were the top models in terms of docking score which required minimum energy to interact with each other (Fig 6, Table 4). *Bph26*-dipeptidyl peptidase IV, *Bph14*-carboxylesterase, *Bph14*-dipeptidyl peptidase IV, *Bph9*-dipeptidyl peptidase IV, *Bph14*-carboxylesterase, *Bph14*-phosphoglycerate kinase 1, *Bph18*-carboxylesterase, *Bph14*-serine protease 6, *Bph26*-phosphoglycerate kinase 1 and *Bph14*-cathepsin L were top 10 models which showed very good interaction with higher hydrogen bonds (Fig 6, Table 4). Most of the salivary elicitors interacted at the LRR region of BPH resistant genes where only few salivary elicitors interacted with the nearby NBS region. *Bph18* showed very poor interaction with the tested salivary elicitors, it may be because it lacks the interactive LRR region (Fig 6). Salivary elicitors of BPH, WBPH and SBPH insects predicted to be interacted with the N terminal LRR region of BPH R genes. Initial LRR repeats of LRR region seems highly interactive with the interactive residues of salivary elicitors (Fig 7).

**Table 4.**
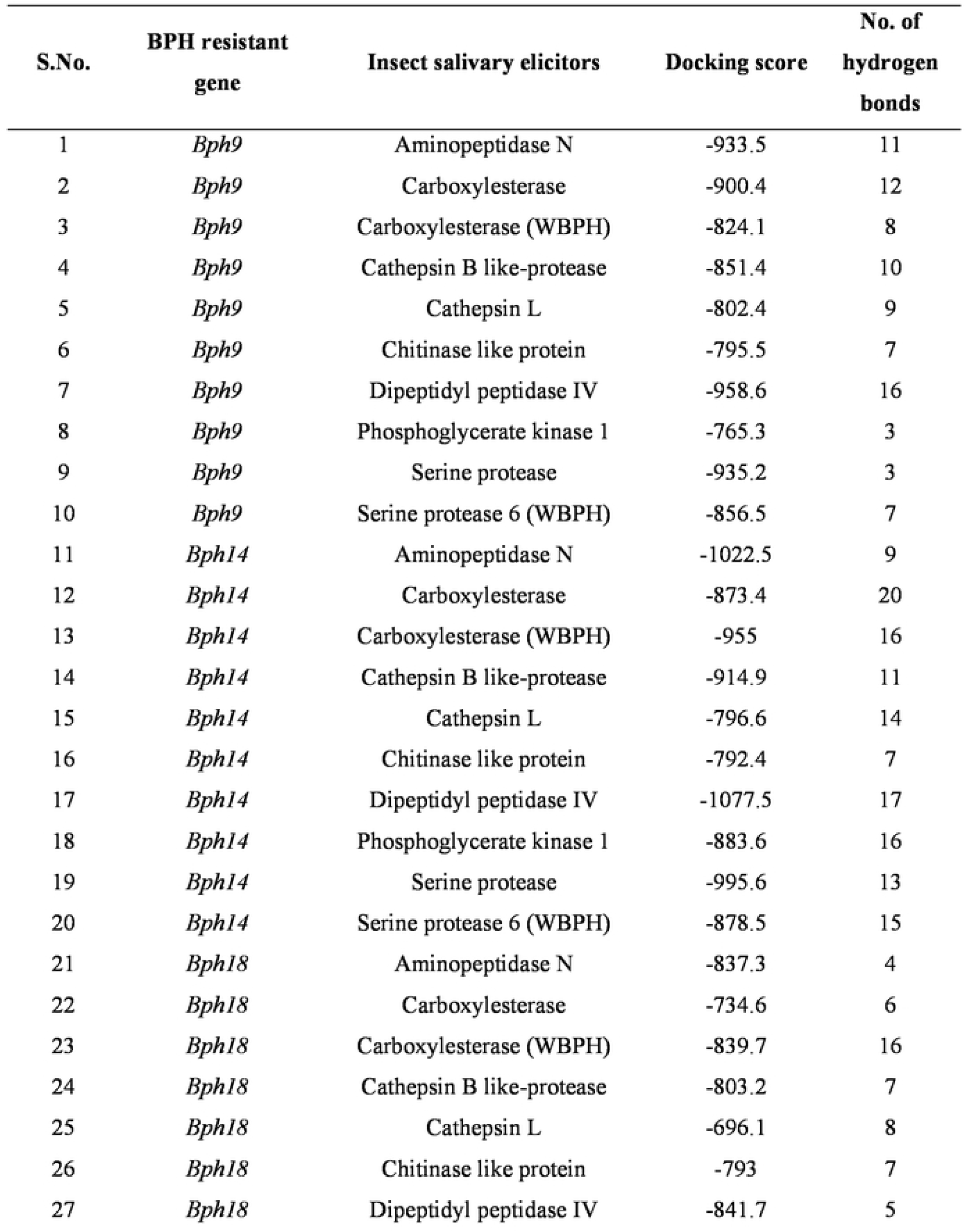
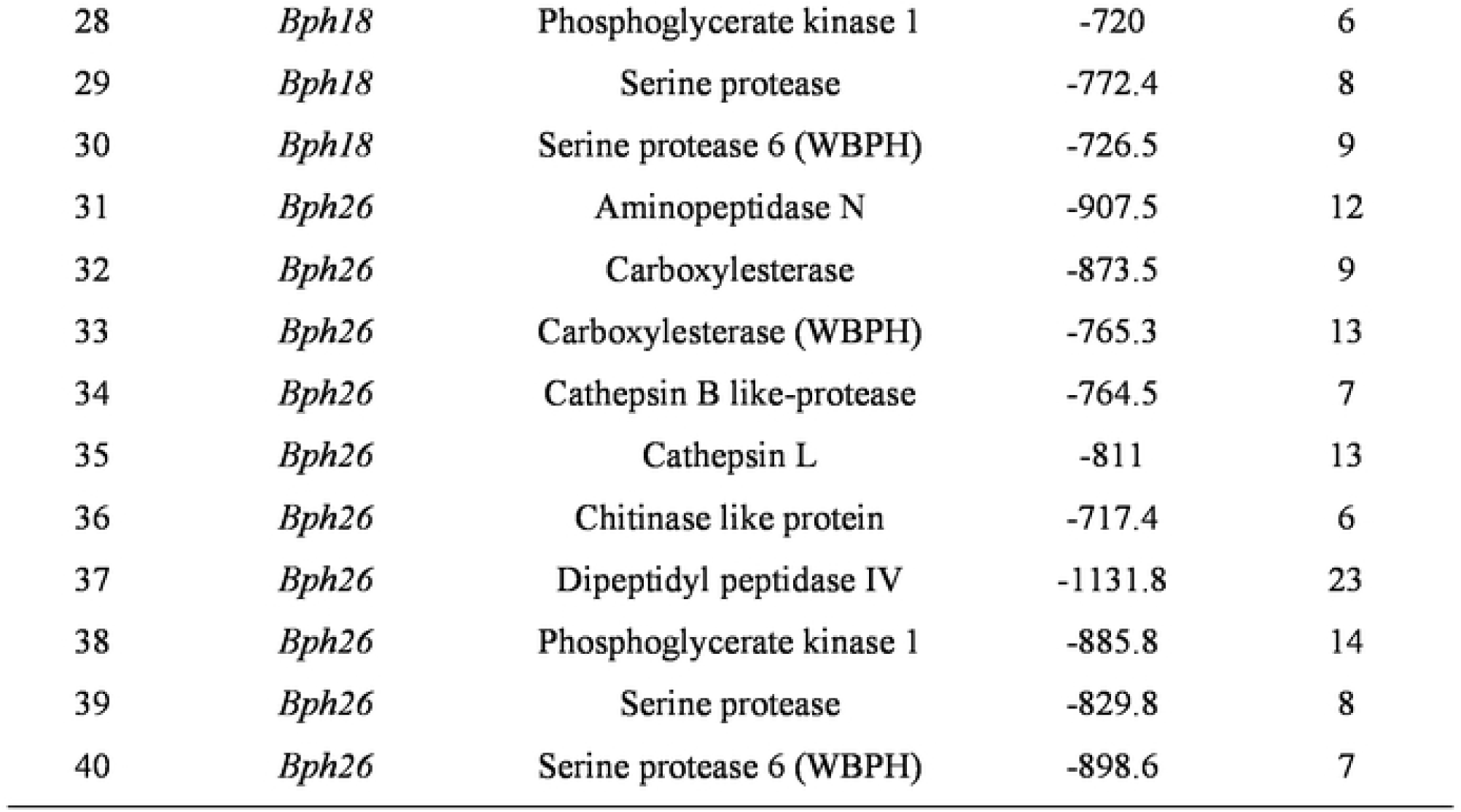
Docking score and hydrogen bond details of BPH resistant R proteins and salivary elicitors of BPH, WBPH and SBPH insects.

| S.No. | BPH resistant gene | Insect salivary elicitors | Docking score | No. of hydrogen bonds |
| --- | --- | --- | --- | --- |
| 1 | *Bph9* | Aminopeptidase N | -933.5 | 11 |
| 2 | *Bph9* | Carboxylesterase | -900.4 | 12 |
| 3 | *Bph9* | Carboxylesterase (WBPH) | -824.1 | 8 |
| 4 | *Bph9* | Cathepsin B like-protease | -851.4 | 10 |
| 5 | *Bph9* | Cathepsin L | -802.4 | 9 |
| 6 | *Bph9* | Chitinase like protein | -795.5 | 7 |
| 7 | *Bph9* | Dipeptidyl peptidase IV | -958.6 | 16 |
| 8 | *Bph9* | Phosphoglycerate kinase 1 | -765.3 | 3 |
| 9 | *Bph9* | Serine protease | -935.2 | 3 |
| 10 | *Bph9* | Serine protease 6 (WBPH) | -856.5 | 7 |
| 11 | *Bph14* | Aminopeptidase N | -1022.5 | 9 |
| 12 | *Bph14* | Carboxylesterase | -873.4 | 20 |
| 13 | *Bph14* | Carboxylesterase (WBPH) | -955 | 16 |
| 14 | *Bph14* | Cathepsin B like-protease | -914.9 | 11 |
| 15 | *Bph14* | Cathepsin L | -796.6 | 14 |
| 16 | *Bph14* | Chitinase like protein | -792.4 | 7 |
| 17 | *Bph14* | Dipeptidyl peptidase IV | -1077.5 | 17 |
| 18 | *Bph14* | Phosphoglycerate kinase 1 | -883.6 | 16 |
| 19 | *Bph14* | Serine protease | -995.6 | 13 |
| 20 | *Bph14* | Serine protease 6 (WBPH) | -878.5 | 15 |
| 21 | *Bph18* | Aminopeptidase N | -837.3 | 4 |
| 22 | *Bph18* | Carboxylesterase | -734.6 | 6 |
| 23 | *Bph18* | Carboxylesterase (WBPH) | -839.7 | 16 |
| 24 | *Bph18* | Cathepsin B like-protease | -803.2 | 7 |
| 25 | *Bph18* | Cathepsin L | -696.1 | 8 |
| 26 | *Bph18* | Chitinase like protein | -793 | 7 |
| 27 | *Bph18* | Dipeptidyl peptidase IV | -841.7 | 5 |
| 28 | *Bph18* | Phosphoglycerate kinase 1 | -720 | 6 |
| 29 | *Bph18* | Serine protease | -772.4 | 8 |
| 30 | *Bph18* | Serine protease 6 (WBPH) | -726.5 | 9 |
| 31 | *Bph26* | Aminopeptidase N | -907.5 | 12 |
| 32 | *Bph26* | Carboxylesterase | -873.5 | 9 |
| 33 | *Bph26* | Carboxylesterase (WBPH) | -765.3 | 13 |
| 34 | *Bph26* | Cathepsin B like-protease | -764.5 | 7 |
| 35 | *Bph26* | Cathepsin L | -811 | 13 |
| 36 | *Bph26* | Chitinase like protein | -717.4 | 6 |
| 37 | *Bph26* | Dipeptidyl peptidase IV | -1131.8 | 23 |
| 38 | *Bph26* | Phosphoglycerate kinase 1 | -885.8 | 14 |
| 39 | *Bph26* | Serine protease | -829.8 | 8 |
| 40 | *Bph26* | Serine protease 6 (WBPH) | -898.6 | 7 |

## Discussion

In *Oryza sativa* (rice), resistant (R) genes (*Bph9, Bph14, Bph18*, and *Bph26*) encoding NBS-LRR proteins have been reported against BPH. These NBS-LRR proteins are approximately 1200 - 1300 amino acids in range (Table 2). BPH specific NBS-LRR are clustered on chromosome 12 (*Bph18, Bph26* and *Bph9*) and chromosome 3 (*Bph14*) along with other NBS clusters (Fig 8). NBS-LRR regions are often clustered in plant genome because of segmental and tandem duplication of chromosomal segments resulting in high gene copy numbers and contributing to the evolution of resistance specificities against insect pests [31–33]. In rice over 500 NBS LRR regions have been identified along its 12 chromosomes of which chromosome 11 comprises 201 loci of NBS gene classes [34]. Similarly, the so far identified BPH resistant genes (39) are mainly clustered in four gene clusters, of which chromosome 12 has the largest cluster reporting 9 of the so far identified BPH resistant loci [35]. Many of the resistant loci identified in this cluster are reported as multi-allelic variants i.e. (*Bph1, Bph2, Bph7, Bph9, Bph10, Bph18, Bph21* and *Bph26* as allelic variants at chromosome 12) and the resistant genes *Bph9, Bph18* and *Bph26* in this cluster encode NBS-LRR proteins [36]. The remaining genes in the cluster i.e. *Bph1, Bph2, Bph7 Bph10, Bph21* have not been functionally characterized. The sequence similarity between *Bph9, Bph18* and *Bph26* was higher confirming them as functional allelic form of BPH resistance. Majority of the NBS-LRR clusters in plants are of similar sequences due to tandem duplication producing closely related NBS genes in the same cluster (e.g., *Bph9, Bph18* and *Bph26* on chromosome 12). However, segmental duplication of chromosomal segments produces related NBS genes located distantly on a chromosome or on a different chromosome (*Bph14* on chromosome 3 highly similar in sequence to *Xa1* and *Xa14* genes on chromosome 4; *Bph9* and *Bph26* on chromosome 12 highly similar in sequence to *Pib* gene on chromosome 1). In spite of sequence similarity, the NBS-LRR genes studied has showed variability in their NBS and LRR regions with amino acid substitutions and deletions (Fig 1A, Fig 1B).

Motif analysis showed the predominant motifs as CC, NBS and LRR in BPH R genes. In *Bph 9* gene, CC domain is able to self-associate and induce cell death phenotype response whereas NBS regulates the activity of CC and LRR region is involved in ligand perception [37]. NBS domain forms a nucleotide binding pocket and act as a molecular switch regulating signal transduction by conformational changes [9]. The CC and NBS domain were highly conserved among *Bph9, Bph18, Bph26* genes and the NBS domain is partially duplicated in the three genes. However, Tamura et al 2016 have cloned and characterized *Bph26* gene as NBS-LRR, but our analysis shows a highly conserved and partially duplicated NBS-I region (Fig 1C) along with NBS-II region in *Bph9, Bph18* and *Bph26* genes (Fig 1A). Domain swap experiments of *Bph9* gene by Wang et al 2021 has showed that multiple intra molecular interaction among CC, NBS and LRR domains are involved in the resistance of *Bph9* gene. The right structural arrangement of the CC-NBSI-NBSII-LRR domains is essential for the induction of cell death phenotype in *Bph9* gene [37]. Recognition of pathogen by LRR region causes a conformational change that is transduced through LRR to NBS II enabling the exchange of ADP for ATP and triggering a second conformational change in the NBS-CC region and further activation of the signalling cascade [38].

LRR domain of NBS-LRR is involved in pathogen effector recognition and the β sheet portion of the LRR region as the possible ligand binding site and contributes to the immune specificity [38]. The presence of effector molecules is recognized by LRR domain triggering a conformational change in NBS domain and thereby initiating the signalling cascade [39]. This LRR region is under diversifying selection with continued evolution against effector/virulence molecules of insect pests. In BPH R genes, the LRR region comprises of an array of β turns with a core of about 26 amino acids containing the Leu-xx-Leu-xx-Leu-x-Leu-xx-Cys/Asn-xx motif (where x is any amino acid) (Fig 1B) Similarly, LRR region was conserved but with varied number of repeats among the BPH genes expect *Bph18* which lacks the LRR region. The solvent exposed residues (x in the consensus sequence) show significantly high ratios of nonsynonymous to synonymous substitutions and are under divergent selection creating variation and evolving of new R genes in a cluster [38,40]. *Bph18, Bph26* and *Bph9* were highly similar in their coiled coil and NBS region with varied number of LRR repeats creating functional alleles in the cluster (Fig 1A, Fig 1B). *In silico* studies confirm that the LRR domains of the BPH proteins are involved in interacting with the elicitor molecules of planthoppers (Fig 7). The LRR arrays of the protein contribute to binding specificity for interacting with variety of ligands providing board spectrum resistance. *Arabidopsis thaliana RPM1* (NBS LRR) disease resistance gene is effective against two avirulent gene products of *Pseudomonas syringae* [41]. However, there are no study that have reported the resistance spectrum of the cloned BPH genes against different planthopper populations expect *Bph14* proved to be effective against WBPH and BPH populations [42]. *Bph18* lacks the LRR region and most of the interacting elicitor molecules were found interacting at the end of the NBS II region (Fig 7). Domain swap experiments on *Bph9 gene* without LRR region resulted in loss of resistance activity against BPH further confirming the ligand recognition by LRR region [37]. *Bph14* with higher number of LRR repeats and variation exhibited higher interaction efficiency with all the tested salivary elicitors providing a broad-spectrum resistance. The number of repeats and variation in the hypervariable region of LRR repeats contribute for new ligand binding specificities providng broader interaction efficiency. The results from the *in-silico* study supports the consideration that plant species adaptation to insect pest elicitors/effector molecules is due to variability that occur in the resistance genes due to continued evolutionary interaction forces between plants and insect pest. Further, insect associated virulent factors that contribute to the resistance have to be studied for better understanding the evolutionary biology of plant insect interaction.

## Conclusion

In the study *in silico* analysis are used for predicting the structure of resistant (R) proteins by comparative modelling and their domains involved in possible interaction with the salivary elicitors of planthoppers. LRR region of BPH R genes were interacting with the elicitor molecules of planthoppers and are the potential lignad binding site. *Bph14* with higher number of LRR repeats exhibited higher binding efficiency with all the tested salivary elicitor molecules. Our *in silico* studies confirms that *Bph14 R* gene resistance protein to be a promising candidate for providing broad spectrum resistance against planthoppers of rice. Further molecular studies can provide better insights into BPH specific R gene-elicitor interaction and activation of signalling cascade against planthoppers of rice.

## Acknowledgement

The corresponding author is acknowledging Sathyabama Institute of Science and Technology for the support. Author K P acknowledging Mr. Carlton Ranjith Wilson Alphonse, Scientific Assistant, Sathyabama Institute of Science and Technology.

---

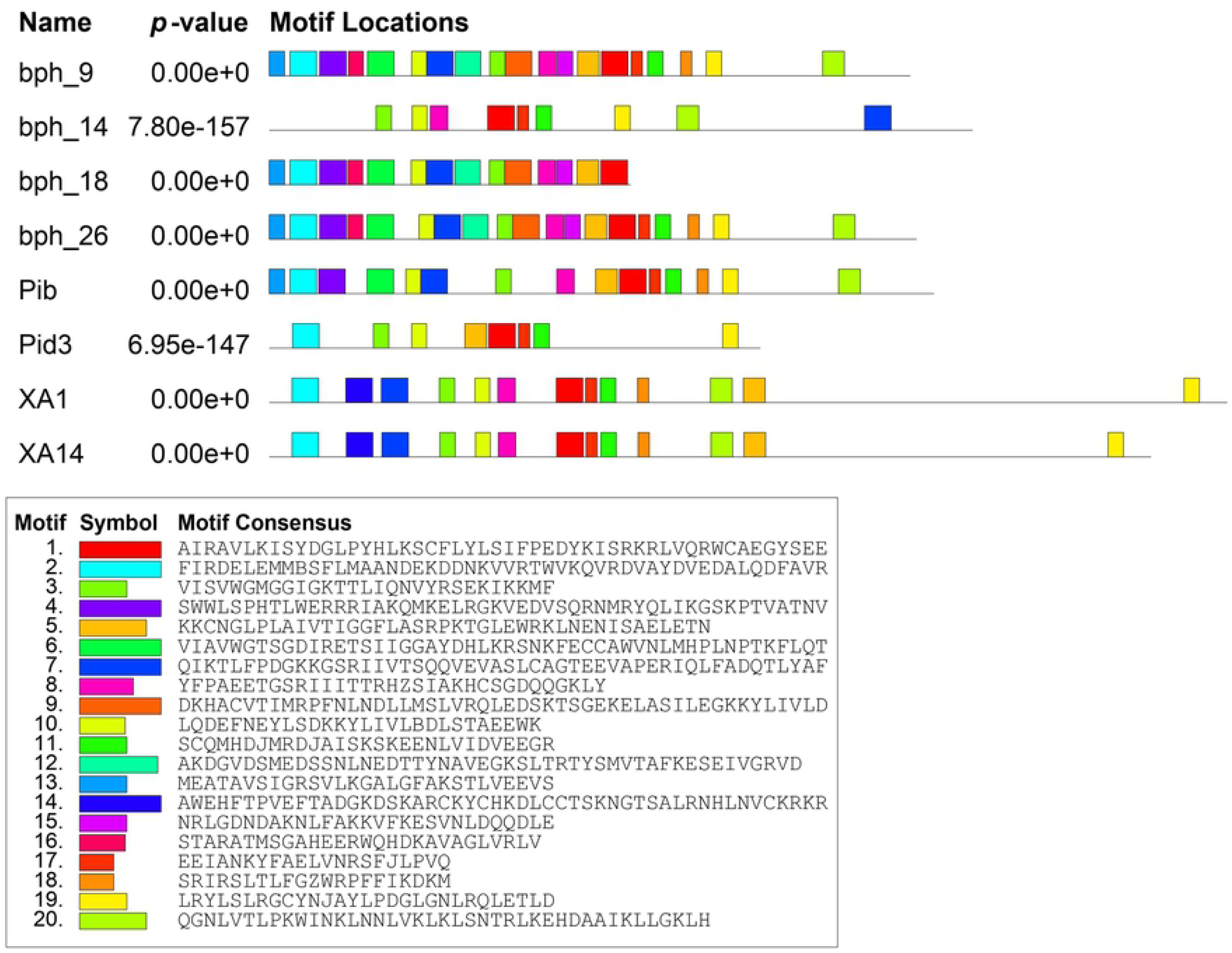

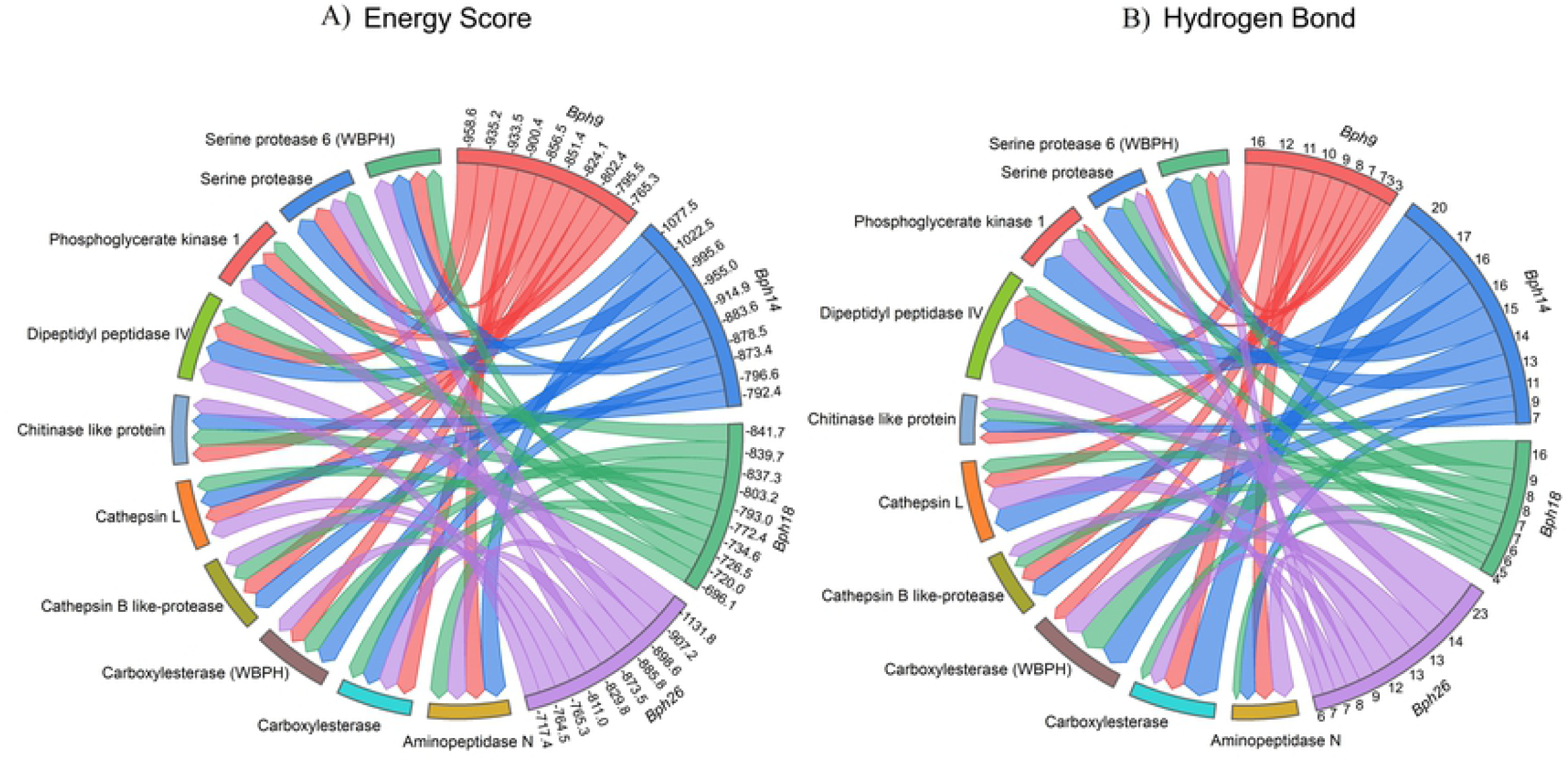

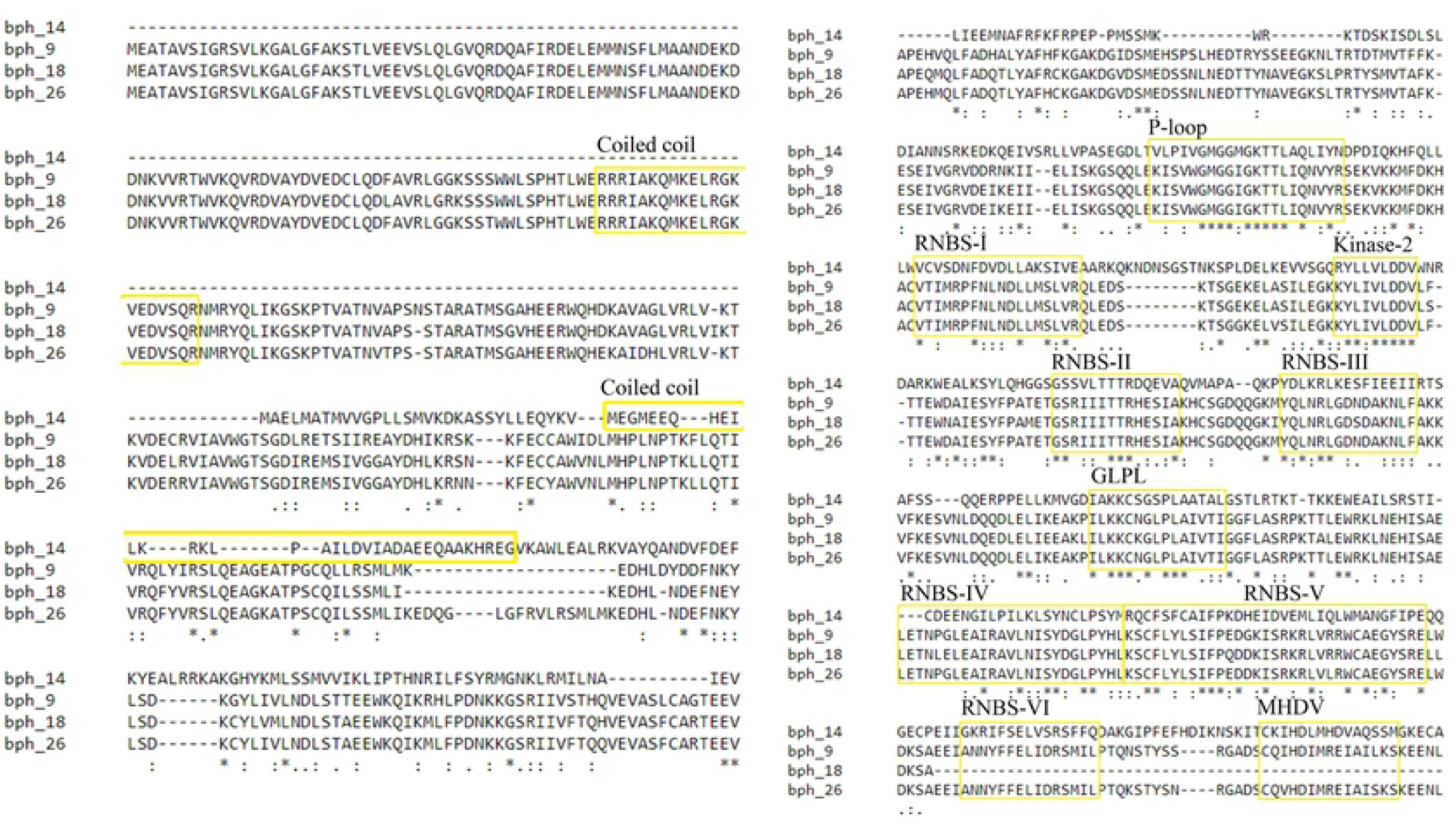

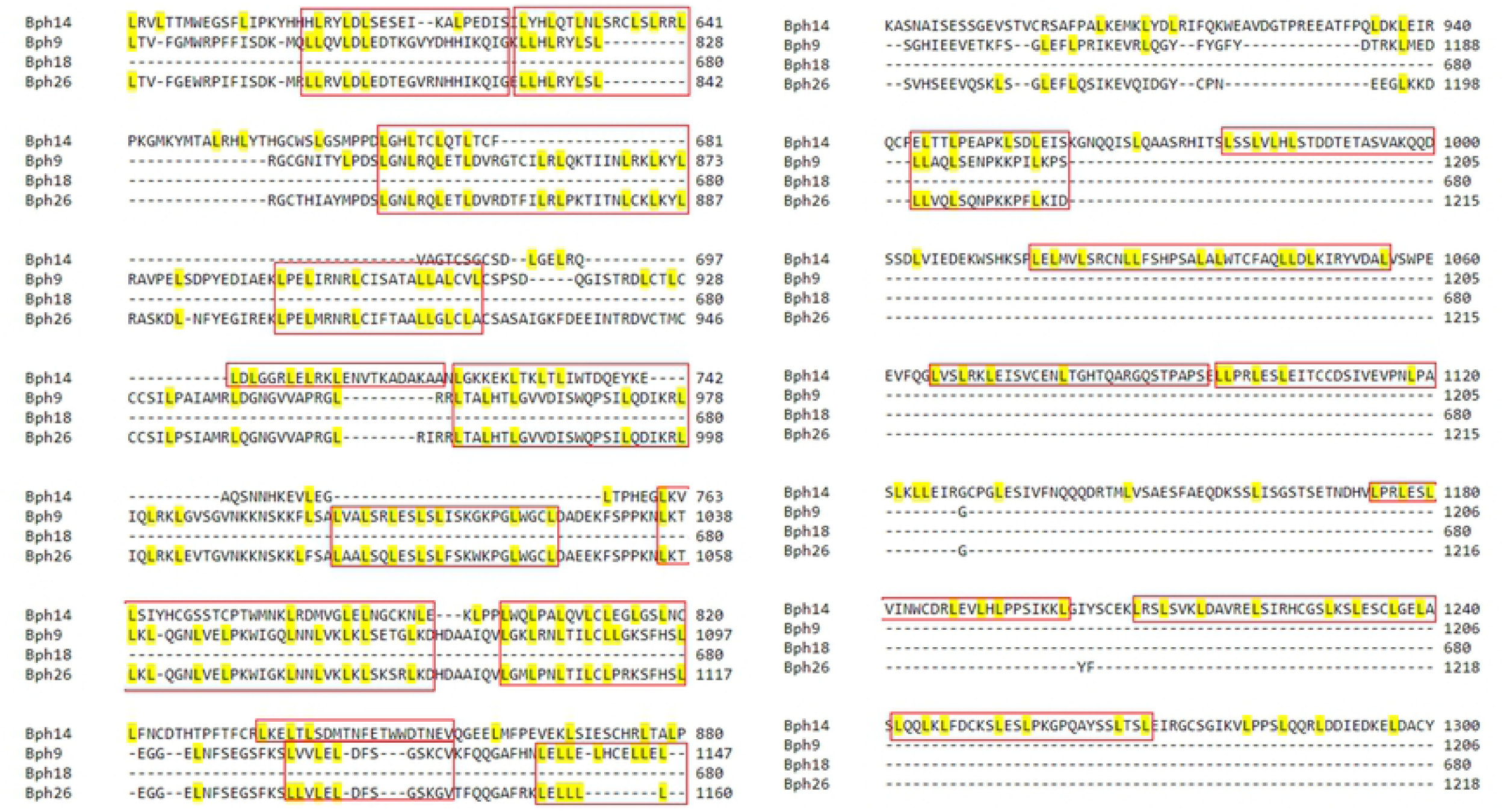

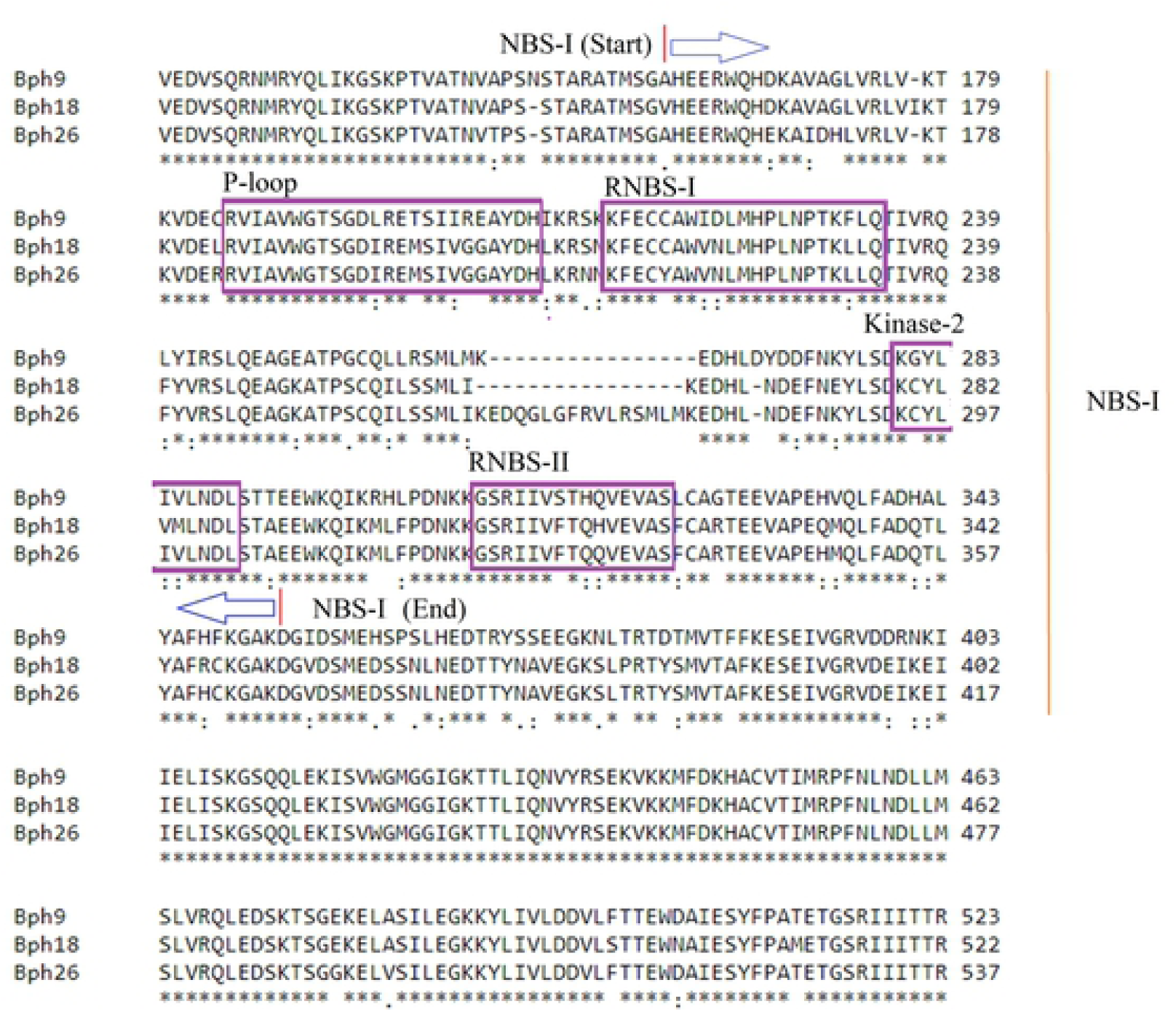

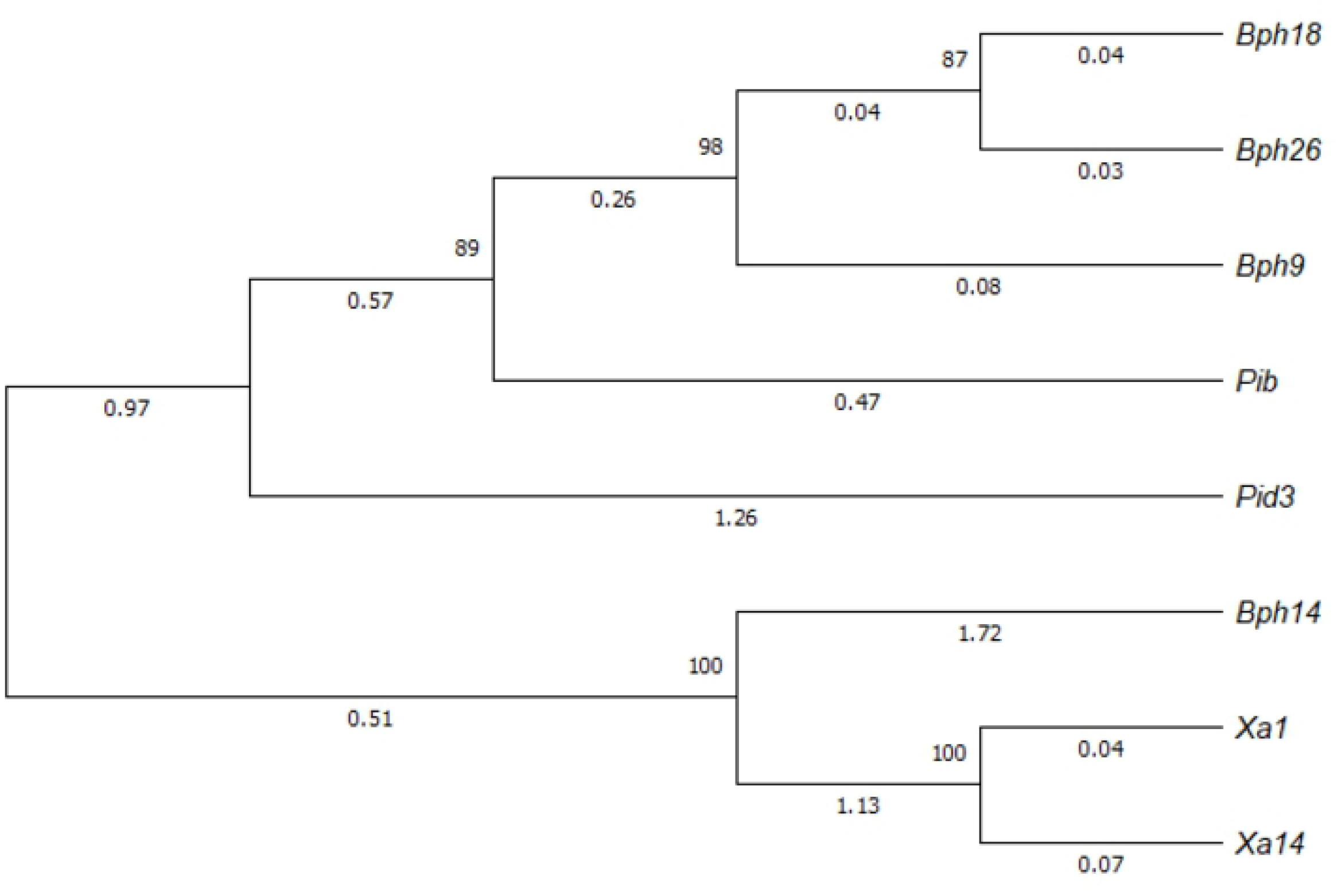

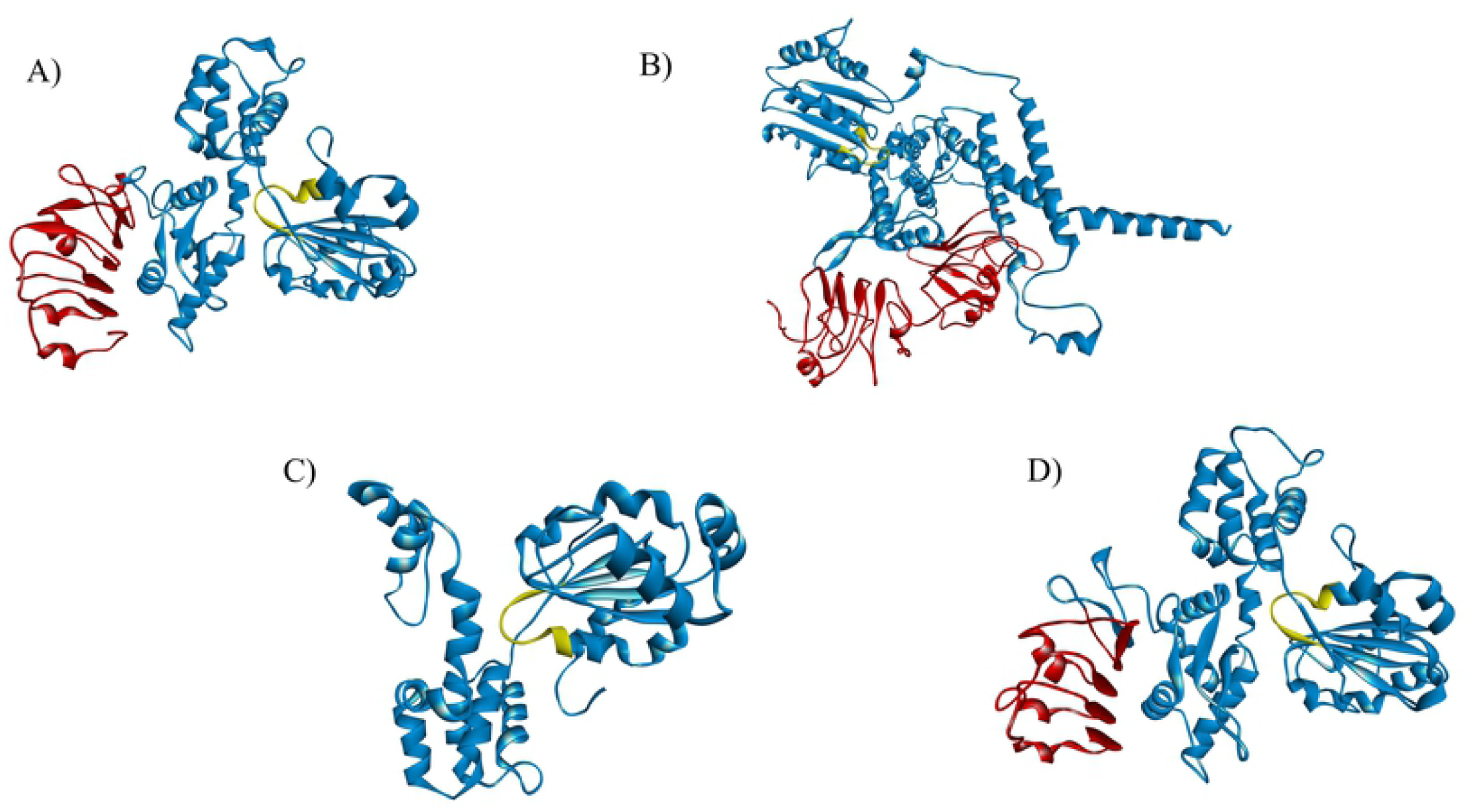

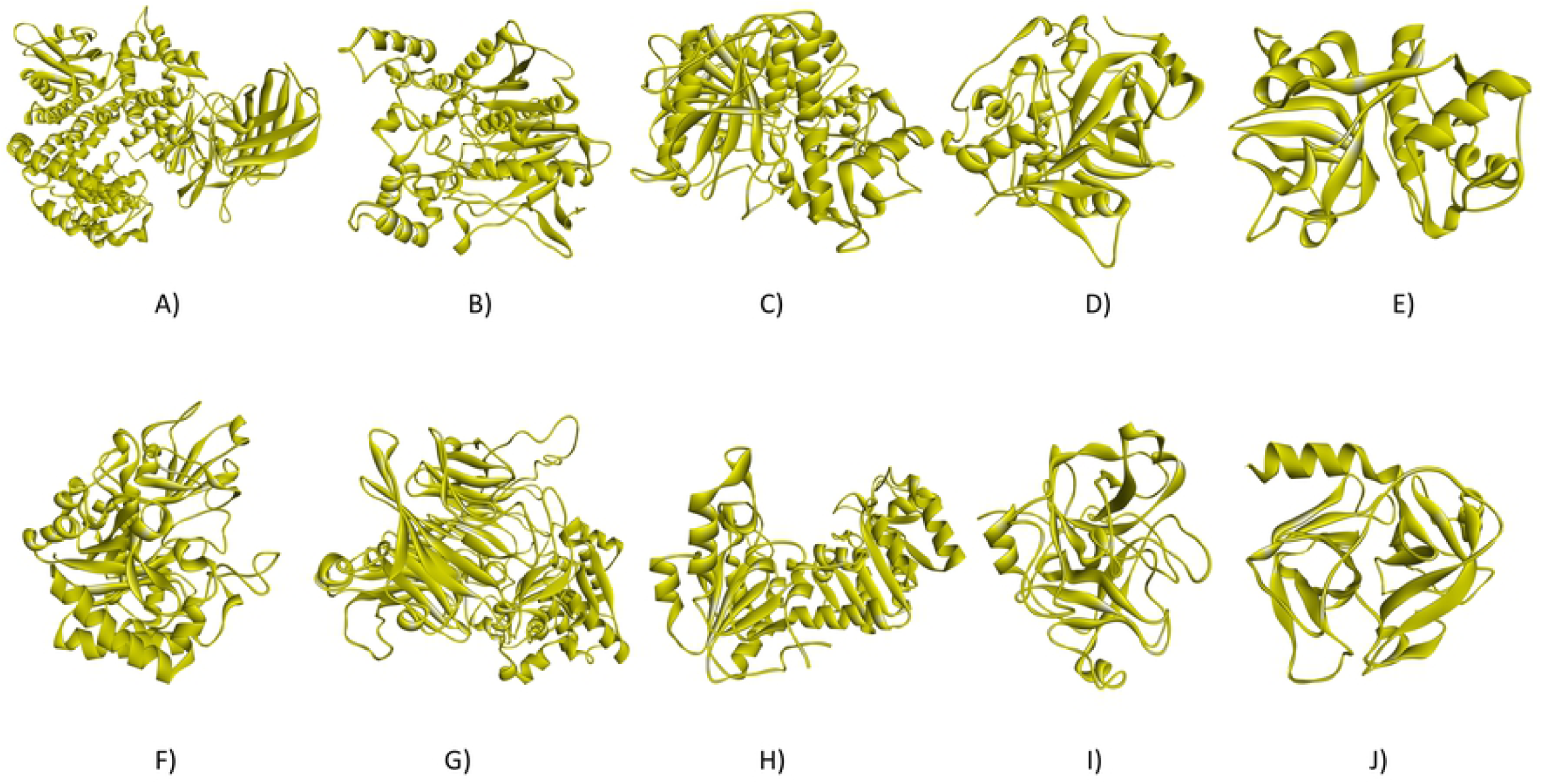

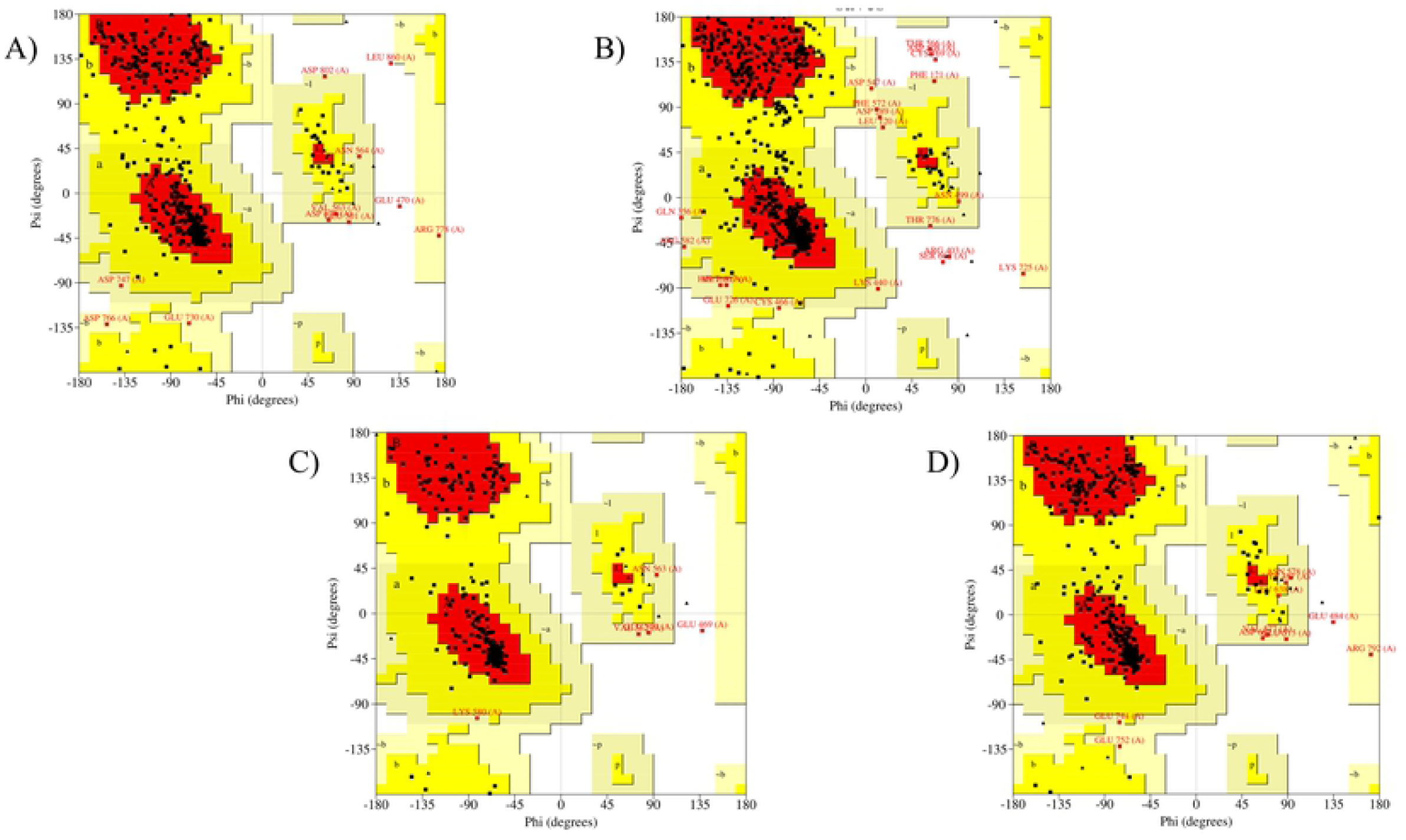

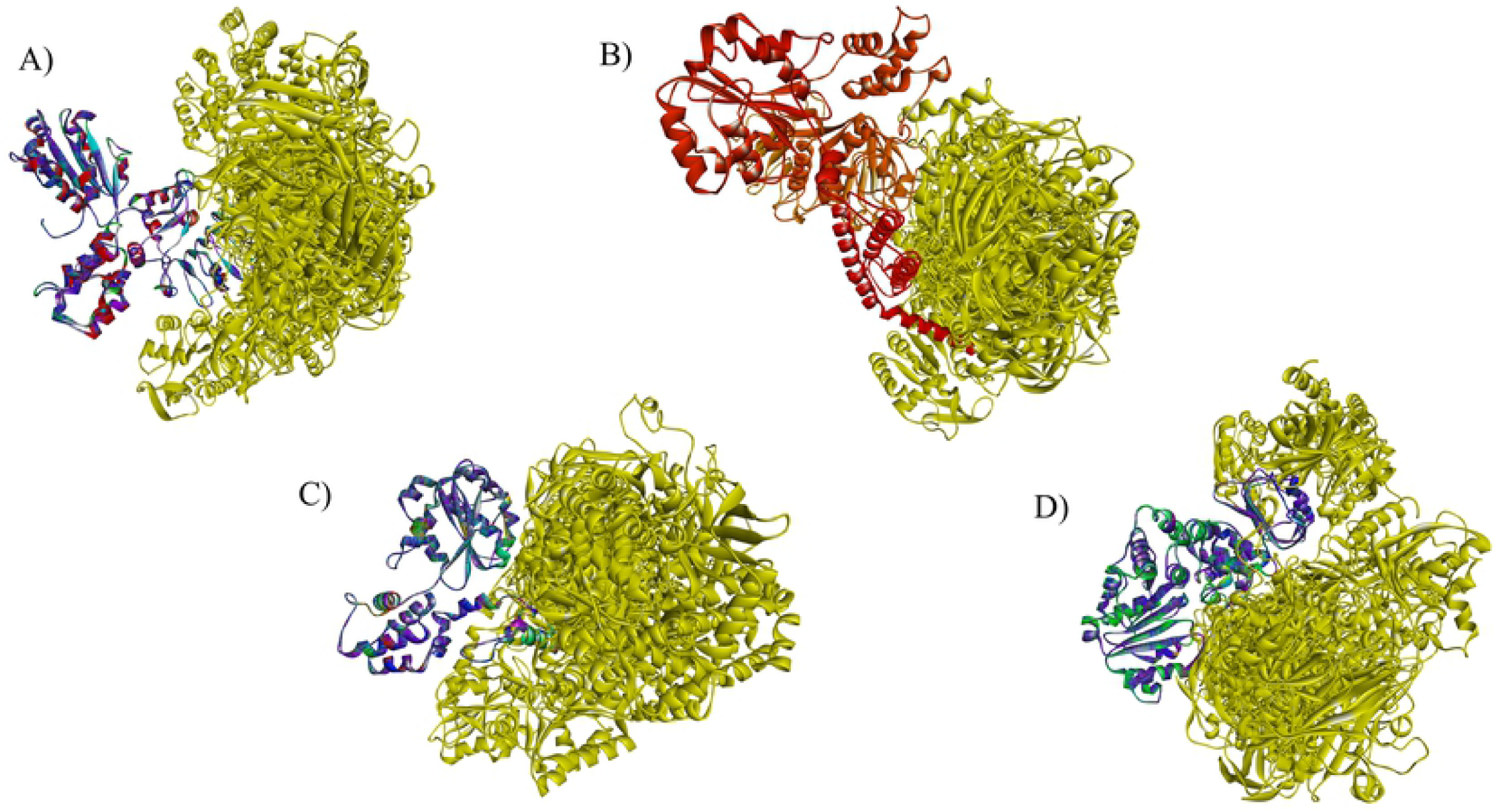

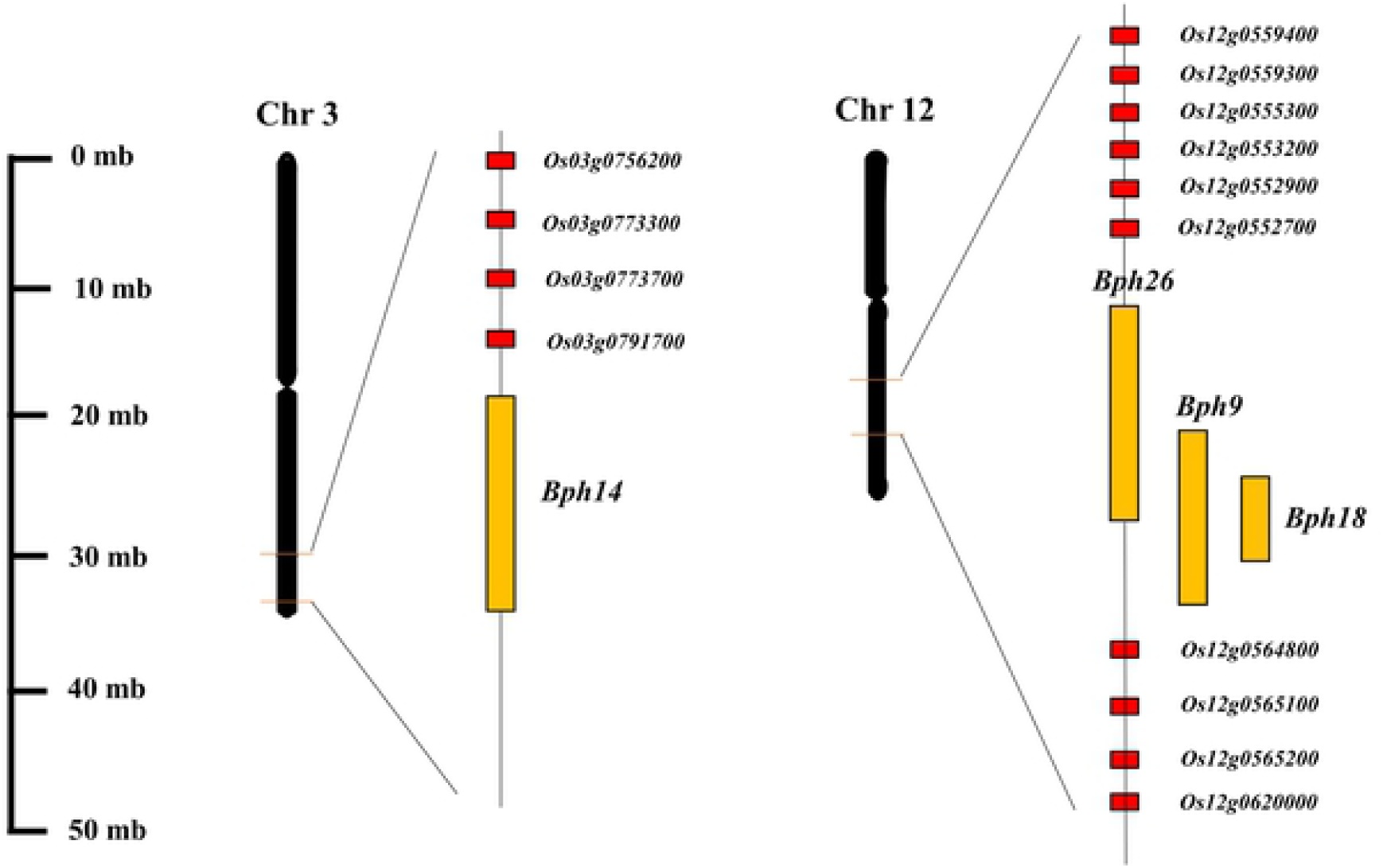

